# ALS-Linked FUS and SOD1 Mutations Elevate MCM2 in Human Motor Neurons

**DOI:** 10.1101/2025.07.09.663892

**Authors:** Kanza Saleem, Andreas Hermann

## Abstract

**Background:** Amyotrophic lateral sclerosis (ALS) is a fatal neurodegenerative disorder characterised by progressive motor neuron loss. In typically post-mitotic neurons, abnormal reactivation of cell cycle regulators and DNA replication licensing factors is observed in ALS pathogenesis. Emerging evidence links components of the minichromosome maintenance (MCM) complex, notably MCM2, to replication stress and genomic instability, implying a mechanistic role in ALS pathogenesis.

**Objective:** To determine whether ALS-associated mutations in FUS and SOD1 influence MCM2 expression and localisation in human induced pluripotent stem cell (hiPSC)-derived spinal motor neurons (MNs).

**Methods:** We differentiated isogenic hiPSC lines carrying FUS P525L-GFP and R495QfsX527, a SOD1 mutant line, and matched wild-type controls into spinal MNs (≥28 days in vitro). QRT-PCR quantified MCM2 mRNA levels (ΔΔCt method; n=3 biological replicates, technical triplicates). Protein expression was assessed by Western blot densitometry (n=3). Subcellular distribution of MCM2 in FUS mutants was evaluated by immunofluorescence (pilot, N=1; ≥50 cells quantified).

**Results:** FUS P525L MNs exhibited a modest, non-significant increase in MCM2 mRNA (1.3-fold vs. WT; *p*=0.1319) but a significant 1.8-fold elevation in MCM2 protein levels (*p*=0.034). The R495QfsX527 line showed a comparable trend at the transcript level (1.2-fold; *p* > 0.05) and a 1.6-fold increase in protein (*p* = 0.041). SOD1 mutant MNs demonstrated a pronounced 2.3-fold MCM2 protein upregulation (*p*=0.008). Immunofluorescence in FUS mutant MNs revealed no significant nuclear-to-cytoplasmic shift in MCM2 localisation, indicating that elevated MCM2 levels are not driven by subcellular mislocalization.

**Conclusion:** ALS-linked FUS and SOD1 mutations upregulate MCM2 protein in human spinal MNs, suggesting post-transcriptional or stability-driven regulation. The absence of relocalisation in FUS mutants shows that this impact is caused by overexpression rather than mislocalisation. MCM2 may be a biomarker of disease-associated replication stress. Future studies will explore whether MCM2 overexpression exacerbates DNA damage or serves as a compensatory response, clarifying its role in ALS pathogenesis.

## INTRODUCTION

The progressive loss of motor neurons in the brain and spinal cord is a hallmark of amyotrophic lateral sclerosis (ALS), also referred to as motor neuron disease. This debilitating neurodegenerative illness causes muscle weakness, paralysis, and eventually respiratory failure (Niedermeyer et al., 2019; Scelsa et al., 2002; Wijesekera & Leigh, 2009). The illness often causes mortality within a few years of diagnosis and affects people all around the world, with a slightly greater frequency in men (Wijesekera & Leigh, 2009; Talbot, 2009). Approximately 10% of cases are familial, although most are sporadic. These instances are associated with genetic mutations in genes such as SOD1, TDP-43, FUS, and C9ORF72 (Hardiman et al., 2017; Wijesekera & Leigh, 2009; Renton et al., 2014). To understand the diagnosis, prognosis, and treatment of ALS, research into new biomarkers and therapeutic targets is still crucial (Swindell et al., 2019). The molecular processes underlying the development of ALS, including DNA damage and poor repair, are becoming increasingly acknowledged. In typically post-mitotic neurons, abnormal reactivation of cell cycle regulators and DNA replication licensing factors is frequently observed (Tokarz et al., 2016). Although neurons are typically post-mitotic, abnormal reactivation of cell cycle machinery, including DNA replication licensing factors, has become increasingly common in ALS. Emerging evidence links components of the minichromosome maintenance (MCM) complex, notably MCM2, to replication stress and genomic instability, implying a mechanistic role in ALS pathogenesis (Zeman & Cimprich, 2014; Fragkos & Naim, 2017). DNA replication licensing protects genomic integrity even under stressful situations by ensuring that DNA is replicated only once per cell cycle, which is controlled by proteins such as the MCM2-7 complex and Cdt1 (Nishitani & Lygerou, 2002). Replication stress and genomic instability can result from the dysregulation of this strictly regulated mechanism, especially when it involves crucial licensing components, such as MCM2. The molecular mechanisms behind DNA damage, inadequate repair, and the onset of ALS are increasingly being recognised. Cell cycle regulators and DNA replication licensing factors are often abnormally reactivated in typically post-mitotic neurons (Tokarz et al., 2016). A member of the minichromosome maintenance complex, MCM2, is essential for DNA replication licensing and cell cycle progression (A.Shetty et al., 2005; Nishitani et al., 2002; Mughal et al., 2019). Dysregulation of this protein has been associated with several malignancies and developmental abnormalities (Blow et al., 2008; A. Shetty et al., 2005; Schmit & Bielinsky, 2021).

Given MCM2’s critical role as a core component of the DNA replication licensing machinery, as well as its role in maintaining genomic stability under cellular stress, studying its expression and regulatory dynamics in the context of ALS, particularly within spinal motor neurons, might offer valuable insights into the molecular mechanisms that underlie motor neuron degeneration. Therefore, the role of MCM2 in ALS pathogenesis, specifically in spinal motor neurons, warrants further investigation to understand its potential contribution to the disease process. This study compares MCM2 expression at both the mRNA and protein levels in hiPSC-derived spinal motor neurons with FUS and SOD1 mutations with those of wild-type controls. Explicitly highlight that, to your knowledge, no one has yet examined MCM2 in ALS patient-derived MNs. Understanding how ALS-associated mutations affect the replication licensing machinery could reveal novel disease pathways.

## MATERIAL AND METHODS

### hiPSC culture, NPCS and Motor neuron differentiation

An isogenic pair of FUS WTGFP and P525L-GFP, 30.1 control iPSCs cell lines with R495QfsX527 cell line and AKC control cell line, along with SOD1 mutant cell line previously reported by Dash et al., 2024; Naumann et al., 2018, were differentiated into MN until at least 28 days in vitro, as described previously (Reinhardt et al., 2013). These were cultured in feeder-free conditions on Matrigel with TeSR-E8 media (StemCell, Germany) and detached with Accutase during maintenance. NPCs and MNs were differentiated from them as described previously (Reinhardt et al., 2013). Briefly, colonies of iPSCs were collected and maintained in media containing 10 µM SB-431542, 1 µM Dorsomorphin, 3 µM CHIR 99021 and 0.5 µM pumorphamine (PMA). After 2 days, the media was replaced with N2B27, which consisted of the factors mentioned above, and DMEM-F12/Neurobasal 50:50 with 1:200 N2 Supplement, 1:100 B27 lacking Vitamin A, and 1% penicillin, streptomycin, and glutamine. On day 4, 150 µM ascorbic acid was added while Dorsomorphin and SB-431542 were withdrawn. On day 6, embryonic bodies were mechanically separated and replated on Matrigel-coated dishes; Matrigel was diluted (1:100) in DMEM-F12 and kept on the dishes overnight at room temperature. The NPCs exhibit a ventralized and caudalized character at this stage, forming homogenous colonies. NPCs were split at a ratio of 1:10–1:20 once a week using Accutase for 10 min at 37° C. Final MN differentiation was induced by treatment with 1 µM PMA in N2B27 exclusively. After 2 days, 1 µM retinoic acid (RA) was added. On day 9, another split step was performed to seed them on a desired cell culture system. To induce neural maturation, developing neurons were treated with N2B27 containing 10 ng/µl BDNF, 500 µM dbcAMP and 10 ng/µl GDNF.

### Mycoplasma testing

The cell lines were regularly checked for mycoplasma testing. We used the Mycoplasma Detection kits for conventional PCR according to the manufacturer’s instructions (Venor GeM, No 11–1025).

### RNA isolation and reverse transcription

RNA was isolated using the Zymo Research Quick-RNA Miniprep Kit (USA). RNA concentration was measured using a Qubit fluorometer or a microplate reader with a NanoQuant plate. For analysis by quantitative polymerase chain reaction (qPCR), RNA samples were converted to cDNA using the High-Capacity cDNA Synthesis Kit (Thermo Fischer Scientific, USA). The reaction was prepared by mixing one µg of RNA with 2 µl of 10X RT Buffer, 0.8 µl of 25X dNTP Mix (100 mM), 2 µl of 10X Random Primers and 1 µl of Reverse Transcriptase. The mix was then filled to the final reaction volume of 20 µl with RNase-free PCR-grade water. In the thermal cycler, samples were incubated at 25 °C for 10 minutes, then at 37 °C for 120 minutes, and boiled at 85 °C for 5 minutes. After cooking, the samples were cooled on ice and stored at -20 °C.

qPCR was performed using the FastStart SYBR Green Master Mix (Roche, Switzerland). The master mix was prepared by mixing 25 µL of FastStart SYBR Green Master, 1.5 µL of 10 µM Primer Forward (final concentration, 300 nM), 1.5 µL of 10 µM Primer Reverse (final concentration, 300 nM), and 17 µL of PCR-grade water. Primer sequences are listed in the table below. Next, 5 µL of cDNA was pipetted into a MicroAmp Reaction Tube, and 45 µL of an appropriate master mix was added. The reaction was performed using either the LightCycler Nano or the LightCycler 480II with the thermal profiler. The qPCR began with a 10-minute initiation step at 95 °C. This was followed by denaturation at 95 °C for 20 seconds and primer hybridisation with extension of the target sequences at 72 °C for 20 seconds. This cycle, without the initiation step, was repeated 40 times and a melting curve was generated at the end. The evaluation of the qPCR is done with the Rotor-Gene Q Series software. For each primer, a concentration test was performed before generating the standard curve to determine the optimal threshold value through internal regression.

Primers are as follows:

GAPDH Forward Primer-CGGAGTCAACGGATTTGGTCGTAT

GAPDH Reverse Primer-ATTGATGACAAGCTTCCCGTTC MCM2

Forward Primer-TTGCTGTAGGGGAACTGACCGA MCM2

Reverse Primer-TGTGCTTGCCACCTGGGTTTTT

### Western blot

Cells were pelleted in ice-cold PBS, and the protein was isolated using RIPA buffer (Thermo Fisher Scientific) supplemented with protease and phosphatase inhibitors (Sigma-Aldrich) after a 40-minute incubation on ice. The samples were then centrifuged at 30,000 × g for 20 min at 4 °C, and the supernatant, containing the total protein fraction, was collected. The protein concentration was quantified through the BCA assay (Thermo Fisher Scientific). SDS-PAGE was performed with the denatured samples containing 20 μg of total protein. The proteins were transferred onto the nitrocellulose membrane (Bio-Rad Laboratories) using the Trans-Blot Turbo Transfer System (Bio-Rad Laboratories). The unspecific binding sites were blocked with 5% skim milk powder (Sigma-Aldrich) in TBS with 0.1% Tween 20 for 1 hour at room temperature with shaking. The membrane was incubated overnight at 4 °C with shaking in TBS containing 3% skim milk and the primary antibody dilution. After washing with TBS 0.1% Tween 20 (3 times for 5 min with shaking), the membrane was incubated with secondary antibodies diluted (1:10,000) for 1 hour at room temperature with shaking. After washing (three times for 5 min with TBS 0.1% Tween 20, and once for 5 min with TBS), the membrane was imaged using an Odyssey 9120 Fluorescent Imager (Li-Cor Biosciences), and the signal was quantified using Image Studio Lite 5.2 (Li-Cor Biosciences). Primary antibodies were used at the following dilutions: MCM2, dilution 1:1000 (ab4461).

### Immunofluorescence and microscopy

Cells were cultivated on the µ-slide 8-well. The cells were fixed with 4% ice-cold paraformaldehyde for 10 min at room temperature. Membranes were permeabilised with 0.2% Triton X-100 in PBS for 10 min at room temperature. The unspecific binding sites were blocked with a blocking buffer (1% BSA, 5% donkey serum, 0.3 M glycine, 0.02% Triton X-100 in PBS) for 1 hour at room temperature. The primary antibodies were diluted to the desired concentration in the blocking buffer and added to the cells, which were then incubated overnight at 4°C. After washing the primary antibody solution with PBS (3 times for 5 min), the secondary antibody dilution (1:500) in blocking buffer was applied, and the cells were incubated for 1 hour at room temperature. Fluorescence microscopy was performed using an LSM 900 confocal microscope (Carl Zeiss) with a 63x oil-immersion lens.

### Quantification and statistical analysis

For all quantitative analyses, at least three independent experiments were performed, based on three different differentiation procedures for neurons or three different passages for stable cell lines. The statistical analysis was implemented using GraphPad Prism 7.0 software. An appropriate statistical test was chosen based on the dataset. The detailed information is provided in each figure legend. All results are reported as mean ± SEM or ±SD. The specific information is provided in each figure legend. ^*^/#p ≤ 0.05; ^**^/##p ≤ 0.01; ^***^/###p ≤ 0.001; ^****^/####p ≤ 0.0001 were considered significant.

## RESULTS

### FUS-ALS mutation upregulates MCM2 Expression in hiPSC-derived Motor Neurons

To investigate whether the FUS-associated ALS mutation alters MCM2 expression, we measured mRNA and protein levels in hiPSC-derived spinal motor neurons. Quantitative real-time PCR demonstrated a modest increase in MCM2 transcript levels in both the FUS P525L and R495QfsX527 mutant lines when compared to their isogenic wild-type counterparts. Although the changes at the mRNA level were not significant, they suggested an upward trend in MCM2 mRNA levels in the mutants (Figure 1A). In contrast, Western blot analysis revealed a strong and consistent increase in MCM2 protein in both mutant strains. The FUS P525L and R495QfsX527 motor neurons exhibited significant increases in protein levels compared to controls, indicating that ALS-associated FUS mutations may primarily influence MCM2 post-transcriptionally or through increased protein stability (Figure 1B). These findings suggest that ALS-associated mutations mainly increase MCM2 expression at the protein level, likely through post-transcriptional regulation or enhanced protein stability.

**FIGURE.**
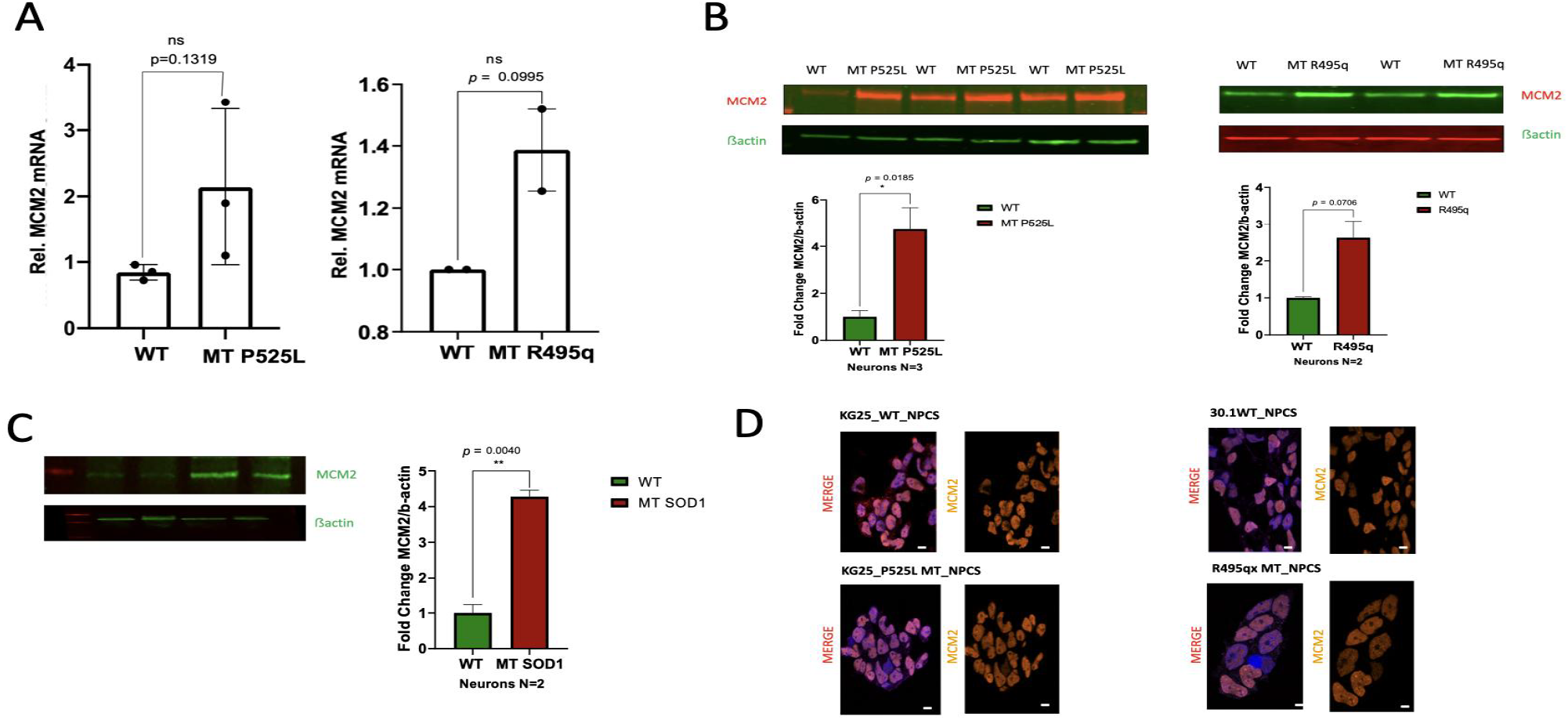
MCM2 expression is elevated in FUS-mutant and SOD1-mutant human motor neurons and NPCs. **(A)** qRT-PCR analysis of MCM2 transcript levels in iPSC-derived motor neurons harbouring the FUS P525L (left) or R495Q (right) mutation compared with their isogenic wild-type (WT) controls. Data are shown as mean ± SEM (n = 3 biological replicates); unpaired two-tailed Student’s *t*-test. **(B)** Immunoblots (top panels) and quantification (bottom panels) of MCM2 protein in FUS-WT versus FUS-P525L (left) or FUS-R495Q (right) motor neurons. Fold-change in MCM2/β-actin is plotted as mean ± SEM (N = 3 for P525L; N = 2 for R495Q); *P = 0.0185 (P525L), P = 0.0706 (R495Q); Student’s t-test. **(C)** Immunoblot (top) and quantification (bottom) of MCM2 protein in MN carrying an ALS-linked SOD1 mutation (MT SOD1) versus their WT. Data are presented as fold-change in MCM2/β-actin (mean ± SD; N = 2); **P = 0.0040 (Student’s t-test). **(D)** Immunofluorescence staining of MCM2 (orange) in neural progenitor cells (NPCs) derived from two independent FUS-WT lines (KG25, 30.1) and their matched P525L or R495Q mutants. Nuclei are counterstained with DAPI (blue) in the merged images (left panels), (N = 1); Scale bar, 10 µm.

### Elevated MCM2 Protein Expression in SOD1-Mutant Motor Neurons

Among the ALS-associated genotypes studied, the most remarkable finding was a significant increase in MCM2 protein in MN with the SOD1 mutant. Compared to wild-type controls, SOD1-mutant spinal motor neurons showed the most tremendous increase in MCM2 protein levels. This significant increase indicates a clear relationship between SOD1 failure and replication licensing dysregulation, implying MCM2 as a potential downstream effector or compensatory marker in response to SOD1-mediated cellular stress (Figure 1C).

### MCM2 Subcellular Localisation Is Unaltered in FUS-Mutant Neural Progenitors

To investigate whether MCM2 dysregulation in ALS is associated with altered subcellular trafficking, we measured the nuclear-to-cytoplasmic distribution of MCM2 in neural progenitor cells generated from FUS-mutant hiPSC lines. Confocal immunofluorescence imaging demonstrated that MCM2 localisation remained predominantly nuclear in both the P525L and R495QfsX527 mutant lines, mimicking the pattern seen in their WT. There was no evident alteration in distribution, suggesting that MCM2 overexpression occurred independently of nuclear export or mislocalization in this cellular setting (Figure 1D).

## DISCUSSION

This study reveals that ALS-linked mutations in FUS and SOD1 lead to a significant increase in MCM2, a key DNA replication licensing factor, in spinal motor neurons. Although MCM2 mRNA levels in FUS-mutant lines increased somewhat but not significantly, protein expression was consistently and significantly elevated. This difference between MCM2 transcript and protein levels suggests post-transcriptional regulation or altered protein turnover, both of which are typically associated with neurodegenerative diseases involving dysfunction of RNA-binding proteins (Naumann et al., 2018). Given that FUS is an RNA-binding protein involved in RNA splicing, stability, and transport, its mutation may affect the post-transcriptional control of transcripts like MCM2. Aberrant FUS might sequester mRNAs into stress granules, hinder typical translation, or change miRNA expression patterns, all of which could have an indirect effect on MCM2 protein levels. Future research that examines MCM2’s connection with ribonucleoprotein complexes or assesses changes in miRNA targeting may reveal these pathways.

Notably, whereas MCM2 dysregulation is widely known in malignancies, where it promotes uncontrolled proliferation, its overexpression in post-mitotic neurons may have quite different implications. Instead of assisting division, high MCM2 levels may cause unscheduled replication licensing, resulting in replication stress and DNA damage responses that eventually lead to neuronal malfunction or death. This contrast demonstrates how, while reactivating cell cycle machinery promotes tumour development, it may also lead to neuronal degeneration.

The SOD1-mutant line exhibited the most significant increase in MCM2 protein, indicating that the replication licensing machinery becomes dysregulated in ALS beyond FUS-driven pathways. The fact that SOD1 mutations are known to cause oxidative stress and DNA damage responses (Dash et al., 2024) suggests that the significant increase in MCM2 might be an attempt to maintain genomic integrity under these stress conditions. This is consistent with MCM2’s recognised involvement in preventing halted replication forks and coordinating replication during stressful settings (Fragkos & Naim, 2017; Zeman & Cimprich, 2014).

Despite the increased protein expression, our immunofluorescence study demonstrated that MCM2 is primarily nuclear in FUS-mutant neural progenitors, with no detectable nuclear-cytoplasmic relocalisation. In contrast to other ALS-related proteins, such as TDP-43 or FUS, our data indicate that subcellular mislocalization is not the primary mechanism underlying MCM2 dysregulation in these animals (Naumann et al., 2018).

Tokarz et al. have reported that abnormal reactivation of cell cycle machinery and DNA replication licensing proteins in post-mitotic neurons is a prevalent hallmark of ALS and other neurodegenerative illnesses (Tokarz et al., 2016). In this context, persistent MCM2 elevation may serve as a compensatory mechanism, helping to mitigate replication stress and DNA damage, or it may represent a maladaptive activation of the cell cycle machinery, exacerbating neuronal dysfunction and cell death (Nishitani & Lygerou, 2002; Blow & Gillespie, 2008). Collectively, our findings reveal MCM2 as a potentially unique molecular marker of replication stress in ALS, underscoring the need for future functional research. MCM2 knockdown, overexpression, or studies on replication fork stability in ALS models may reveal whether its dysregulation causes neurodegeneration or merely reflects downstream cellular stress responses.

## LIMITATIONS

This work utilised a limited number of ALS-mutant hiPSC lines, which may not have captured the entire variability of the disease. The subcellular localisation analysis of MCM2 was conducted as a pilot study with a limited sample size (N = 1), which limited the robustness of the findings. Functional experiments were not used to determine the direct effect of MCM2 overexpression on neuronal viability or DNA damage response. Furthermore, the findings are limited to in vitro systems and require confirmation in patient tissue or in vivo ALS models for greater translational value.

